# Bioelectrical signaling via domain wall migration

**DOI:** 10.1101/570440

**Authors:** Harold M. McNamara, Rajath Salegame, Ziad Al Tanoury, Haitan Xu, Shahinoor Begum, Gloria Ortiz, Olivier Pourquie, Adam E. Cohen

**Affiliations:** Department of Physics, Harvard University; Harvard-MIT Division of Health Sciences and Technology; Department of Chemistry and Chemical Biology, Harvard University; Department of Genetics, Harvard Medical School; Department of Pathology, Brigham and Women’s Hospital; Department of Chemistry, University of California Berkeley; Howard Hughes Medical Institute; School of Physics, Peking University

## Abstract

Electrical signaling in biology is typically associated with action potentials, transient spikes in membrane voltage that return to baseline. The Hodgkin-Huxley equations of electrophysiology belong to a more general class of reaction-diffusion equations which could, in principle, support patterns of membrane voltage which are stable in time but structured in space. Here we show theoretically and experimentally that homogeneous or nearly homogeneous tissues can undergo spontaneous spatial symmetry breaking into domains with different resting potentials, separated by stable bioelectrical domain walls. Transitions from one resting potential to another can occur through long-range migration of these domain walls. We map bioelectrical domain wall motion using all-optical electrophysiology in an engineered stable cell line and in human iPSC-derived myoblasts. Bioelectrical domain wall migration may occur during embryonic development and during physiological signaling processes in polarized tissues. These results demonstrate a novel form of bioelectrical pattern formation and long-range signaling.

## Introduction

In 1952, Hodgkin and Huxley introduced a mathematical model of action potential propagation in the squid giant axon, based on nonlinear dynamics of electrically coupled ion channels.^1^ In the same year, Alan Turing proposed a model for biological pattern formation, based on diffusion and nonlinear reaction dynamics of chemical morphogens.^2^ These two seemingly unrelated models have an underlying mathematical kinship: both are nonlinear reaction-diffusion equations, first order in time and second order in space. Thus, from a mathematical perspective, one expects parallel classes of solutions. These solutions can be organized by whether they are uniform or patterned in space, and stable or varying in time (Fig. 1). All four combinations of spatial and temporal structure have been observed in chemical reaction-diffusion systems ^3^, but only three of the four have been reported in the field of electrophysiology. We thus sought to observe the fourth class of electrophysiological dynamics: spontaneous spatial symmetry breaking in a nominally homogeneous tissue.

**Figure 1.**
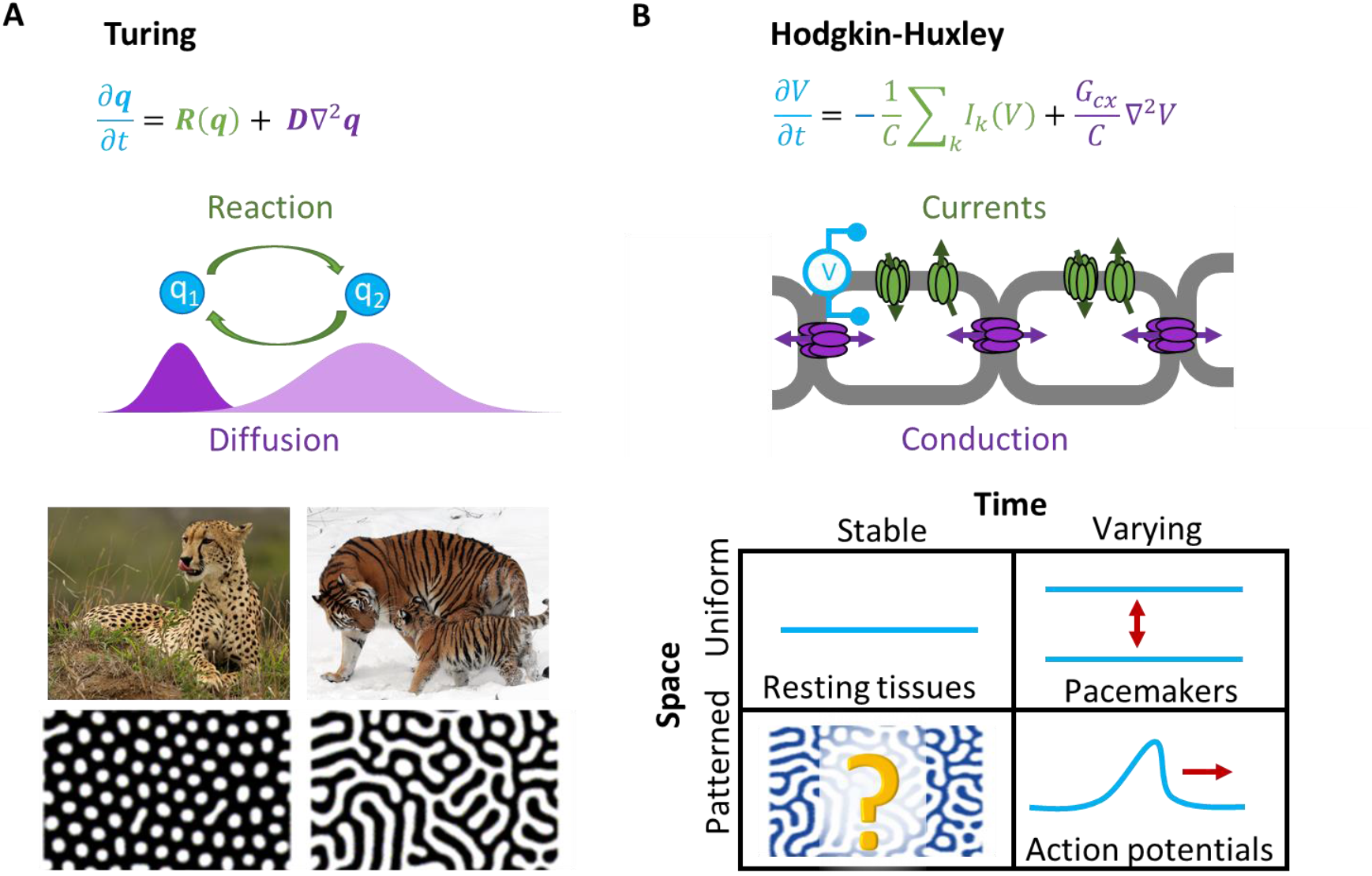
Biochemical and bioelectrical pattern formation. A) In chemical Turing patterns, a nonlinear chemical reaction coupled to diffusion leads to spontaneous formation of stable concentration patterns like ones seen in Nature. Here ***q*** = (q_1_, q_2_) is the vector of reagent concentrations, ***R***(***q***) is the nonlinear relation between concentration and reaction rate, and ***D*** is the vector of diffusion coefficients. B) The Hodgkin-Huxley equation has the same structure as the Turing equation. Here *V* is the membrane voltage, *C* is the membrane capacitance, *I*_k_ is the current through the k^th^ ion channel, and *G*_cx_ is the connexin conductivity. The chart shows possible solutions to the Hodgkin-Huxley equation, classified by variation in space and time. Spatially varying but temporally constant patterns are a little-explored possibility in electrophysiology. Images of natural and simulated patterns adapted from Wikipedia and Kondo et al^3^.

Spatial symmetry breaking might emerge during slow transitions in membrane potential, such as occur during embryonic development and in signaling processes in peripheral organs. While pattern-forming processes in electrophysiology have been contemplated,^4–7^ unambiguous observations with clear mechanistic interpretations have been lacking. Part of the experimental challenge comes from the difficulty of spatially mapping membrane voltage. Patch clamp measurements of membrane potential probe the voltage at only discrete points in space, and are thus ill-suited to mapping spatial structure. Recent advances in voltage imaging facilitate spatially resolved measurements ^8, 9^, and optogenetic stimulation offers the prospect to tune the electrophysiological state of a tissue and perhaps to drive it into a regime of spontaneous symmetry breaking.

Here, we explore these ideas experimentally in engineered cells expressing the inward-rectifying potassium channel K_ir_2.1 and the channelrhodopsin CheRiff. While this two-component cellular model is so simple as to appear almost trivial, we find that coupled ensembles of these cells show richly diverse transitional behaviors, including electrical bistability, bioelectrical domain walls, and noise-induced breakup into discrete electrical domains. We further show that similar dynamics occur in human induced pluripotent stem cell (iPSC)-derived myocytes during differentiation. Our results demonstrate bioelectrical pattern formation and domain wall motion as generic mechanisms by which tissues can switch from one membrane voltage to another.

### Bistable membrane voltages

The lipid bilayer cell membrane behaves, electrically, as a parallel plate capacitor. Transmembrane protein channels can pass ionic currents which alter the intracellular charge, and hence the membrane voltage. In a single cell or a small isolated patch of tissue, the membrane voltage follows:

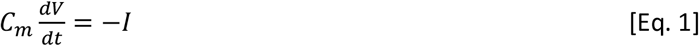

where *C_m_* is the membrane capacitance and *I* is the current through all ion channels (outward positive). Channel gating dynamics can impose a nonlinear and history-dependent relation between *I* and *V* which causes complex dynamics in excitable cells.

Resting potential 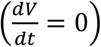 in most polarized cells is set by an inward rectifier potassium channel, K_ir_. The current through the K_ir_ channel is *I_Kir_* = *g*_*K*_*x*_∞_(*V*)(*V* − *E*_*K*_), where *g*_K_ is the conductance (proportional to the number of channels in the membrane), and *x*_∞_(*V*) captures the voltage-dependent gating of the channel.^21^ The term (*V* − *E*_*K*_), where *E_K_* ∼ −90 mV is the potassium Nernst potential, accounts for the electrochemical driving force for ions to cross the membrane. The function *I_K_ir__*(*V*) crosses the x-axis at the potassium reversal potential. Inward rectification implies a drop in K_ir_ current at more positive potentials. Together these attributes give K_ir_ channels a non-monotonic *I*-*V* relationship as in Fig. 2a.^4, 16^ To a good approximation, the K_ir_ conductance depends on present voltage only, not on history.

**Figure 2.**
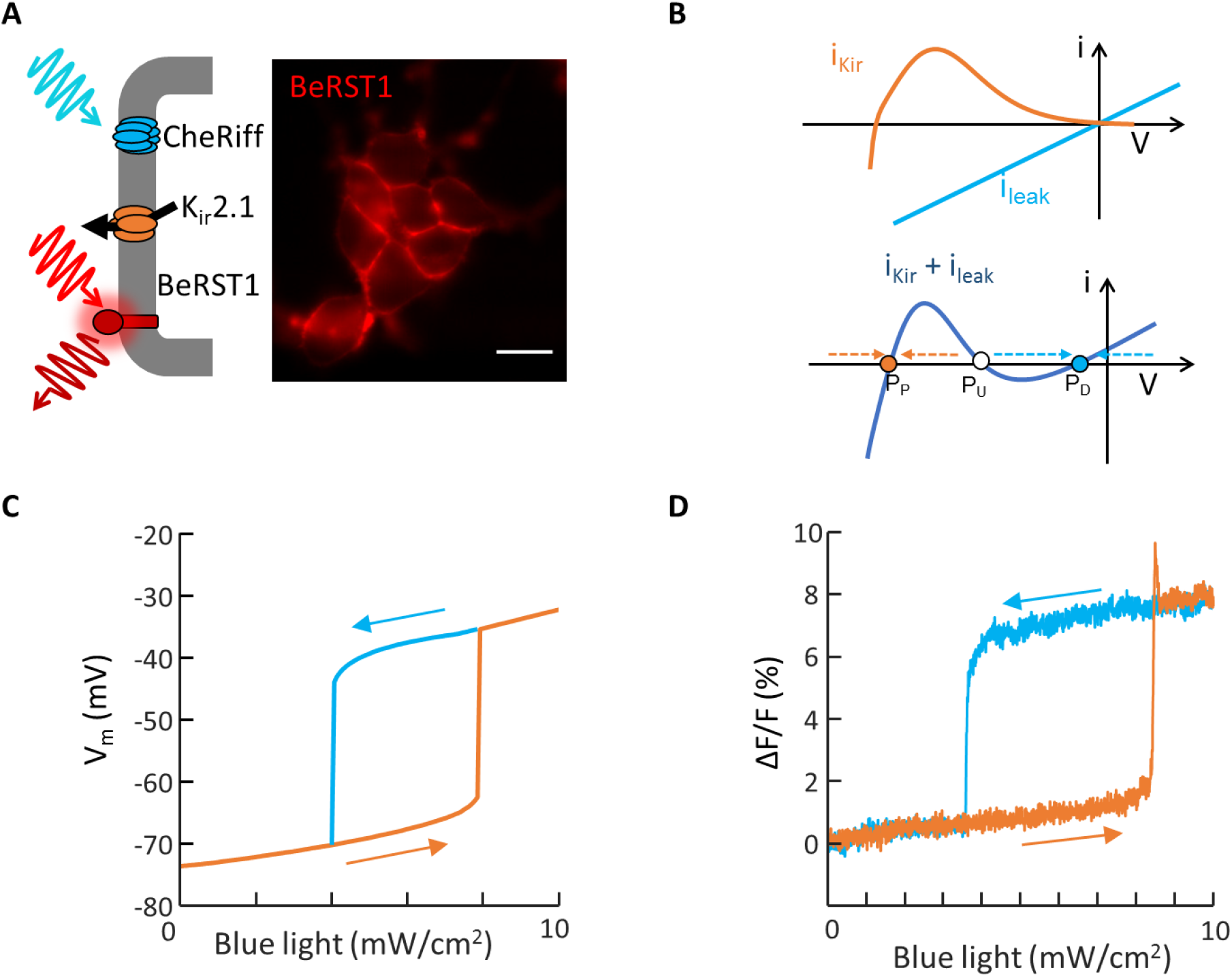
Electrophysiological bistability in an engineered cell line. A) Bi-HEK cells expressed an inward-rectifier potassium channel, K_ir_2.1, and a light-gated ion channel, CheRiff. Fluorescence image of bi-HEK cells labeled with the fluorescent voltage-sensitive dye BeRST1. Scale bar 20 μm. B) Expression of an inward rectifier potassium channel (e.g. K_ir_ 2.1) and a non-selective leak conductance (e.g. channelrhodopsin) are sufficient, together, to produce electrical bistability. C) Numerical simulations showing hysteresis of steady-state membrane potential under ramped optogenetic stimulation. Simulations were based on measured *I*-*V* curves of bi-HEK cells (*Methods*). D) Optical measurements of membrane voltage in a small cluster of bi-HEKs, under ramped optogenetic stimulation.

Cells typically have one or more leak conductances. We consider the simplest case: an Ohmic leak with reversal potential 0 mV and conductance *g*_l_, leading to a straight line *I-V* relation, *I*_leak_ = *g*_l_ *V*. Leak conductances can be gated by external variables, e.g. by chemical ligands or mechanical forces. Below we use a non-selective cation-conducting channelrhodopsin, CheRiff, as a leak conductance where the value of *g*_l_ is readily tunable via blue light ^8^.

The total current is the sum of the K_ir_ and leak currents (Fig. 2b). When *g_l_* dominates, one has a single depolarized fixed point (*I* = 0) near 0 mV (P_D_). When *g*_K_ dominates, one has a single polarized fixed point near −90 mV (P_P_). When *g_l_* and *g*_K_ are approximately balanced, one has an N-shaped *I*-*V* curve which crosses the x-axis three times. This situation implies coexistence of stable fixed points P_D_ and P_P_ with an unstable fixed point (P_U_) in between, leading to overall bistability.^17, 18^ From a dynamical systems perspective, this situation is analogous to the bistability observed in the famous *E. coli* Lac operon system.^19, 20^

We genetically engineered a HEK293 cell line that stably expressed K_ir_2.1 and CheRiff (Fig. 2d, *Methods*).^22, 23^ We call these bistable-HEK cells (bi-HEKs). Patch clamp measurements on small clusters (∼50 μm diameter) of bi-HEKs revealed a non-monotonic *I*-*V* curve, which could be driven through two saddle-node bifurcations by light (Supplementary Fig. 1). We performed numerical simulations of a cell governed by Eq. 1, using the measured *I*-*V* curve (*Methods*). Under continuous variation in blue light the membrane potential underwent sudden jumps at saddle node bifurcations where P_U_ annihilated either P_D_ or P_P_. The jumps occurred at different values of blue light in the polarizing and depolarizing directions, leading to hysteresis (Fig. 2c), i.e. within the hysteretic region, the membrane voltage was bistable. We used a far-red voltage-sensitive dye, BeRST1 ^24^, to report the membrane voltage in small clusters of bi-HEKs as homogeneous blue illumination was slowly increased (0 to 10 mW/cm^2^ over 25 s) and then decreased at the same rate. The membrane voltage showed abrupt transitions and hysteresis, in close agreement with the numerical simulations (Fig. 2d).

Thus, cells expressing K_ir_ + leak exhibited a form of non-genetic electrophysiological memory: the steady-state membrane voltage was not uniquely specified by the ion channels alone. Rather, in the hysteretic regime the steady-state voltage depended on the history of ionic currents, which could in turn depend on the history of stimuli to the cell or, in principle, on the history of gene expression (e.g. whether the leak or the K_ir_ channel was expressed first).^4^

### Bioeletrical domains in extended tissues

In an extended tissue, neighboring cells can be coupled by gap junctions. When the voltage on a cell deviates from the mean of its neighbors, ionic currents flow through the gap junctions to minimize this deviation. The dynamics then become:

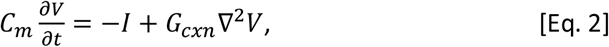

where G_cxn_ is the sheet conductance due to gap junction channels. When the membrane potential is bistable (i.e. the ratio *g*_l_/*g*_K_ is in the hysteretic portion of Fig. 2f), different regions of the tissue may sit at different resting potentials, P_U_ and P_D_.^25^ A domain wall then emerges at the interface between these regions (Fig. 3a). Dimensional analysis implies that the domain wall width scales as 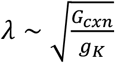.

**Figure 3.**
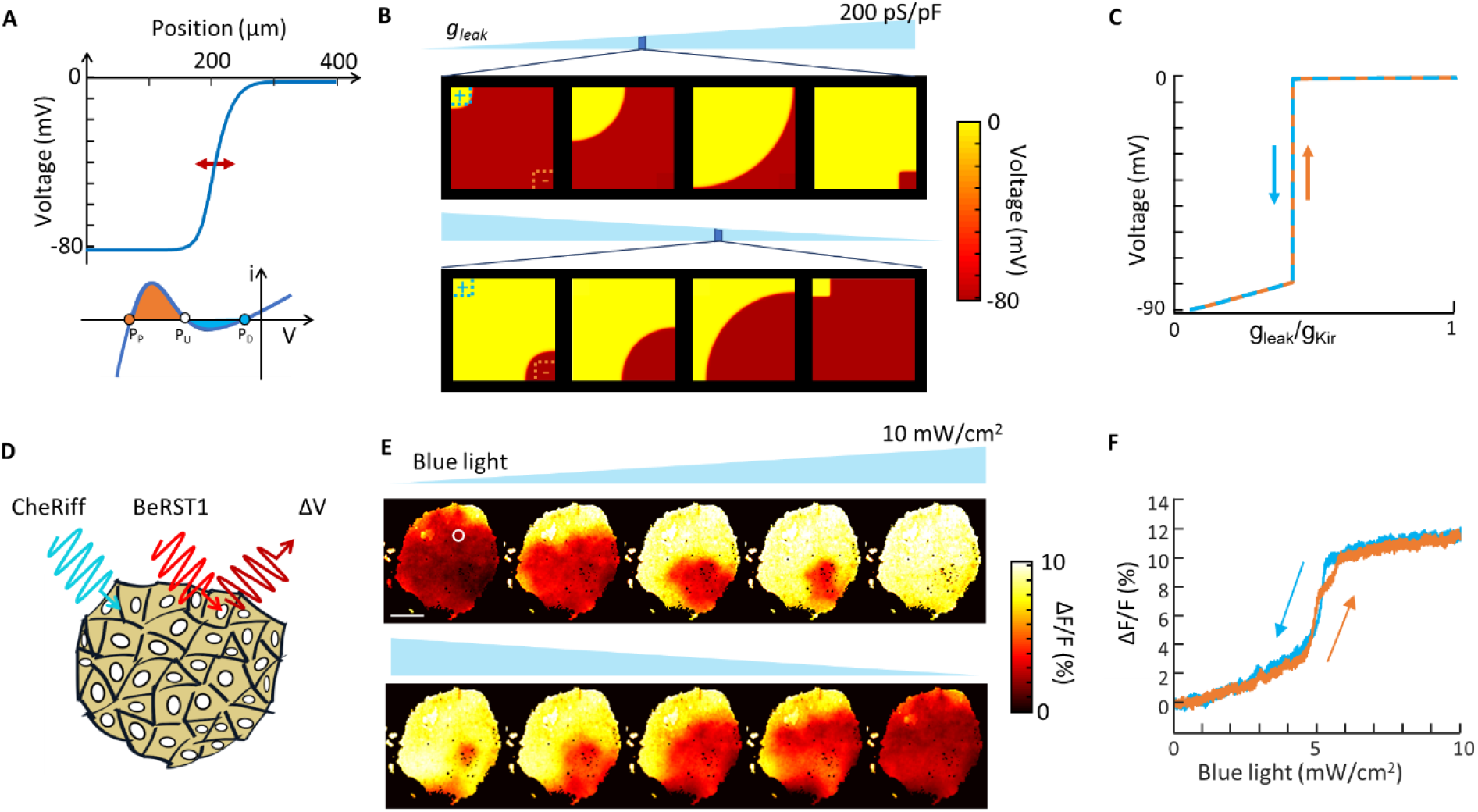
Bioelectric domain walls in an engineered cell line. A) Top: Profile of a bioelectrical domain wall in one dimension, calculated via numerical solution for a balanced K_ir_ and leak current. Bottom: in a homogeneous tissue, the domain wall migrates in a direction set by the relative areas of the orange and blue shaded regions of the *I*-*V* curve, favoring the fixed point with the larger shaded region. B) Simulation of domain wall growth in a homogeneous tissue with two discrete defects to nucleate transitions (clamped at V = 0 on the top left, clamped at V = −90 mV on the bottom right). The transitions in the bulk tissue occur over a narrow range of *g*_leak_. See Supplementary Movie S1. C) Simulation of membrane voltage in the tissue in (B) as a function of leak conductance. D) Confluent islands of bi-HEK cells were illuminated with uniform blue light to stimulate CheRiff, and with red light to elicit voltage-dependent fluorescence of BeRST1. E) Fluorescence images of an island of bi-HEKs under gradually increasing optogenetic stimulation. Scale bar 1 mm. See Supplementary Movies S2 and S3. F) Fluorescence as a function of optogenetic stimulus strength from the region circled in white in (E). Domain wall migration led to a step-like change in membrane potential without hysteresis.

Numerical simulations of Eq. 2 yielded the domain wall structure shown in Fig. 3a (*Methods*). In a homogeneous tissue, the domain wall remained stationary only when the K_ir_ and leak conductances were perfectly balanced. Otherwise the domain wall migrated to expand the territory of the stronger conductance (Supplementary Fig. 2).^26^ In two dimensions, simulations predicted that bioelectrical domains nucleated at defects (e.g. cells that expressed only leak or only K_ir_) and spread through the tissue (Fig. 3b, Supplementary Movie S1). In a tissue that was homogeneous but for an arbitrarily low density of nucleation points, the hysteresis vanished and the transition between depolarized and polarized states was abrupt (Fig. 3c). Thus the collective nature of the transition in an extended tissue was predicted to convert a gradual change in ionic currents into a highly sensitive phase change-like switch in membrane potential.

HEK cells endogenously express connexins 43 and 45 which mediate nearest-neighbor electrical coupling,^27, 28^ so we reasoned that confluent monolayers of bi-HEK cells might support bioelectrical domain walls. We performed optogenetic stimulation and voltage imaging experiments in confluent islands of bi-HEKs with dimensions ∼2 × 2 mm, corresponding to ∼4×10^4^ cells (Fig. 3d). Initially (in the absence of optogenetic stimulation) the tissue was homogeneously polarized. Illumination with dim blue light led to nucleation of depolarized domains near the tissue boundaries. As the blue light further increased, the domain walls migrated across the tissue, until the whole tissue was depolarized (Fig. 3e, Supplementary Movies S2 and S3). The fluorescence intensity of most regions in the island showed step-like depolarization with little hysteresis, consistent with the theoretical predictions. These experiments demonstrated quasi-static domain walls in a nominally homogeneous tissue, a hallmark of spontaneous electrophysiological symmetry breaking.

No tissue is perfectly uniform, so we explored via simulation tissues with cell-to-cell variations in expression of K_ir_ or leak (*Methods*). Noisy ion channel expression introduced an effective friction for domain wall motion, stabilizing droplet-like domains of high or low voltage and broadening the transition in tissue-average voltage under a ramp in *g*_l_ (Supplementary Fig. 3). Sufficiently strong heterogeneity led to stick/slip saltatory domain wall motion. The tissue-average voltage then showed Barkhausen-like fine-structure noise (Supplementary Fig. 4). Tissue heterogeneity also restored some degree of hysteresis in the tissue-average voltage, and, when strong enough, broke the tissue into discrete domains that switched independently. The predicted voltage dynamics of coupled cells expressing leak + K_ir_ thus exhibited many of the features found in magnetization of a disordered soft ferromagnet.^29^

Signatures of noisy ion channel expression were observable in our experiments on island cultures. Due to regional variations in the depolarizing transition point, the voltage averaged over the whole island showed a broader transition than did local measurements (Supplementary Fig. 4). The island-average voltage showed Barkhausen-like fine-structure noise, and a small amount of hysteresis, indicative of stick-slip domain motion. Numerical simulations of islands with noisy gene expression recapitulated these effects (Supplementary Fig. 4). In cultures where transient transfection of K_ir_2.1 led to enhanced cell-to-cell variability in *g*_K_ (*Methods*), we observed breakup into regions in which the voltage showed hysteretic zero-dimension-like behavior and regions which showed smooth and reversible depolarization, in concordance with simulations (Supplementary Fig. 4, Supplementary Movies S4 and S5).

### Electrical bistability and hysteresis during myogenesis

Early embryonic tissue has a membrane voltage, V_m_ ∼ 0 mV ^10^. During myogenesis, myoblast precursors polarize electrically, exit the cell cycle, and fuse into myocytes whose resting potential is ∼-85 mV.^15^ Expression of K_ir_2.1 initiates this hyperpolarization.^30^ In mammals, myoblast precursors couple transiently via gap junctions during differentiation and prior to fusion.^31^ We thus hypothesized that bistability and bioelectric domain wall motion might occur during myogenesis.

We performed all-optical electrophysiology experiments in human induced pluripotent stem cell (hiPSC) derived myoblasts as they differentiated into myocyte fibers *in vitro* (Fig. 4a). HiPSC myoblasts were seeded at low density, lentivirally transduced to expresses CheRiff, and then allowed to proliferate to form a confluent monolayer (Methods; Fig. 4a). The cells were then differentiated into myocytes. After one week of differentiation, cells expressed stained positive for myogenin, PAX7, and myosin heavy chain, and adopted an elongated fiber-like morphology, indicative of differentiation toward mature myocytes (Fig. 4b). RNAseq measurements on matched samples showed a significant increase in K_ir_2.1 expression during the differentiation process (5.6-fold, *p* <0.001) and high expression of gap junction proteins Cx43 and Cx45 throughout (Methods). We performed voltage imaging under ramped wide-field optogenetic stimulation at two time-points during differentiation to test for signatures of electrical bistability in isolated cells and domain wall motion in confluent cultures.

**Figure 4:**
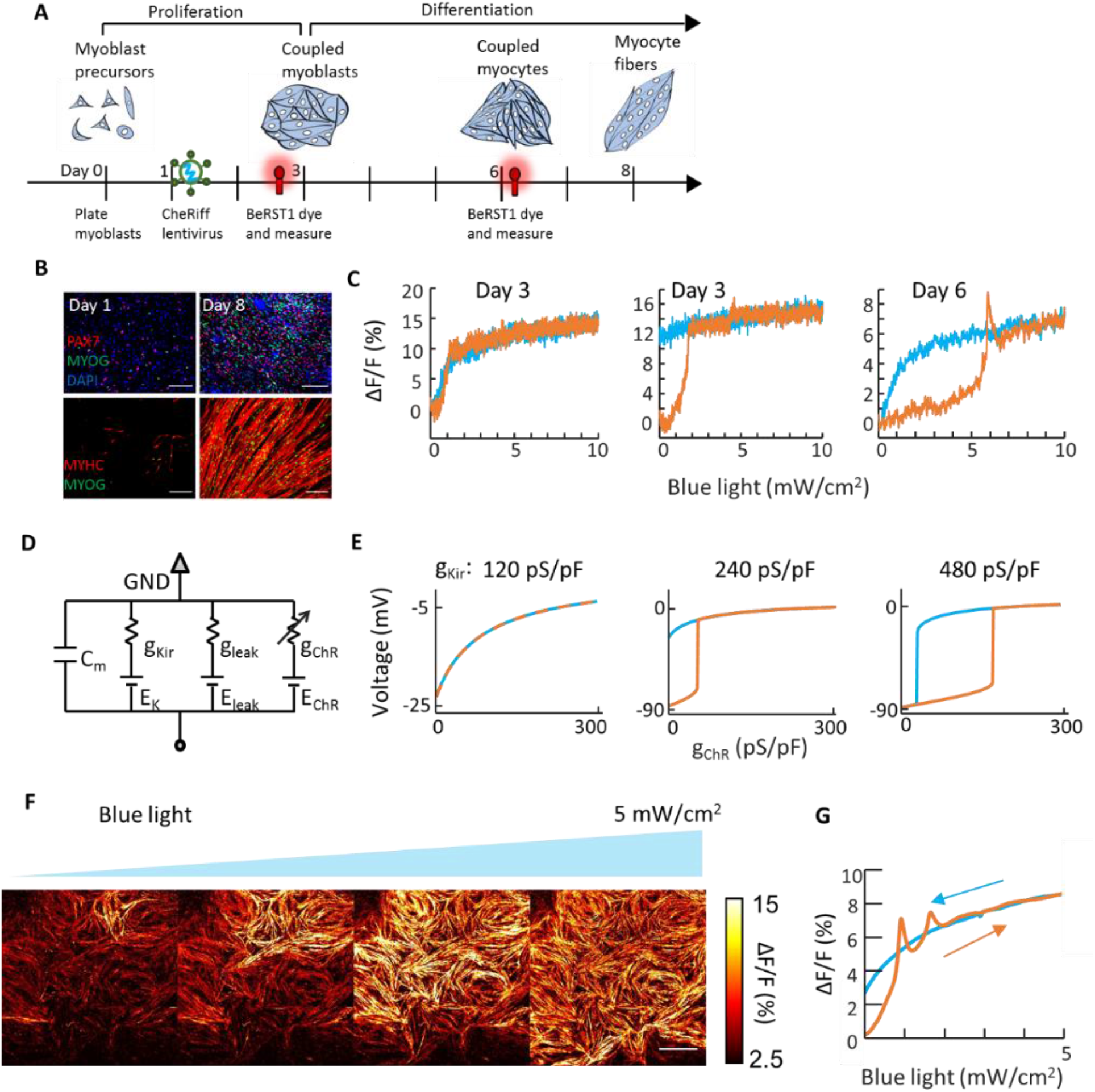
Bioelectric domain wall propagation in stem cell-derived myocytes. A) Timeline for differentiation, viral transduction, and measurement. B) Immunocytochemistry staining of myocyte cultures during differentiation. Stains show PAX7, myogenin (MYOG), and myosin heavy chain (MYHC). Scale bars 200 μm. C) Optical measurements of membrane potential in individual isolated myoblasts at different times after differentiation. D) Simple electrical circuit model for myoblasts. The K_ir_ channel was modeled as a nonlinear conductance, the leak was treated as Ohmic with E_leak_ = −20 mV, and the channelrhodopsin was treated as Ohmic with E_ChR_ = 0 mV. E) Simulations of optogenetically induced changes in membrane voltage at different values of g_Kir_. All other parameters of the simulation were kept constant between the three panels. F) Bioelectrical domain wall migration in a monolayer of electrically coupled myocytes (measured 3 days after start of differentiation). Scale bar 1 mm. See Supplementary Movie S6. G) Optical measurements of membrane voltage as a function of optogenetic stimulus strength in the confluent culture. Depolarization-activated inward currents led to spikes atop the optogenetically induced depolarizations.

In myoblast precursors that had not yet reached confluence (day 3), we observed heterogeneous responses to ramped optogenetic stimulation: Cells showed either a smooth response with saturation-like behavior and little hysteresis (67%, 34 of 51, Fig. 4c left) or a step-wise depolarization, which did not reverse upon cessation of the optogenetic stimulus (33%, 17 of 51, Fig. 4c middle). In immature myocytes mechanically dissociated from a confluent culture (day 6, 3 days after start of differentiation), we observed sub-populations with behavior similar to day 3 (smooth depolarization with no hysteresis: 47%, 42 of 89; step-wise, irreversible depolarization: 29%, 26 of 89). We also observed a new sub-population comprising cells with closed hysteresis loops that resembled the bi-HEKs (24%, 21 of 89, Fig. 4c right, Supplementary Fig. 5, *Methods*).

These three seemingly disparate behaviors could all be explained by a simple model containing a leak, a channelrhodopsin, and K_ir_ expression which increased on average between day 3 and 6 (Fig. 4d, S4). At the lowest K_ir_ level, the *I*-*V* curve was monotonically increasing, so channelrhodopsin activation shifted a single voltage fixed point in the along the *I* = 0 axis. This led to a continuous, and reversible, change in voltage (Fig. 4e). At intermediate K_ir_ level, the *I-V* curve was N-shaped and crossed the *I*=0 axis three times in the absence of channelrhodopsin activation. Blue light drove step-wise depolarization via a saddle node bifurcation. The depolarized state remained stable in the absence of optogenetic drive, leading to non-recovering depolarization. At the highest K_ir_ level, the hysteresis curve shifted to the right and the cells repolarized in the absence of optogenetic drive. Thus, a simple model with one tuning parameter captured the three qualitatively distinct single-cell responses to channelrhodopsin activation.

In confluent monolayers at day 6, we used patterned optogenetic stimulation to excite a portion of the tissue. The evoked action potentials propagated beyond the stimulated region, confirming the presence of gap junctional electrical coupling (Supplementary Fig. 6). Under spatially homogeneous ramped optogenetic stimulation, we observed optogenetically induced domain wall propagation (Fig. 4f, Supplementary Movie S6). The presence of domain wall propagation was surprising, considering that only a minority (24%) of the isolated cells were bistable. Simulations showed that the due to strong electrotonic coupling, the global behavior of a tissue could be dominated by a minority of cells expressing K_ir_2.1 (Supplementary Fig. S2). As in the bi-HEKs, the whole-tissue average voltage showed little hysteresis as a function of optogenetic drive (Fig. 4g), consistent with depolarization via domain wall migration. These observations show that differentiating myoblasts exhibit electrical bistability when isolated and collective domain wall migration during an essential step of myogenesis.

In contrast to the bi-HEK cells, the myoblasts also supported propagation of regenerative action potential waves. These waves manifested as spikes in the whole-tissue fluorescence during a gradual optogenetic depolarization (Fig. 4g). The additional depolarizing drive associated with these spikes caused the waves to propagate rapidly across the tissue, without disruption from the defects which could pin the motion of domain walls.

## Discussion

*In vitro*, muscle cells must be aggregated to differentiate, a phenomenon called the “community effect”.^32^ Our results show that electrical coupling can mediate community effects, i.e. that the collective electrical dynamics of coupled cells can be strikingly different from the individuals, even if all cells are identical. Domain wall migration mediates polarization in extended tissues, whereas isolated cells or small clumps must polarize all at once. Consequently, under ramped K_ir_2.1 expression, an electrically coupled, extended tissue will polarize before an isolated cell or small patch, even if all other conditions are identical. K_ir_2.1 expression is required for the expression of the myogenic transcription factors Myogenin (MyoG) and Myocyte Enhancer Factor-2 (MEF2).^30^ Disruption of gap junction coupling is sufficient to disrupt myogenesis.^33^ Together, these observations suggest that bioelectric domain walls might play a functionally important role in myogenesis.

Gap junctions are necessary for proper formation of many tissues during development, including in heart, liver, skin, hair, cartilage, bone, and kidney,^34, 35^ though the physiological roles of these gap junctions remain unclear. Our work suggests that gap junction-mediated bioelectrical domain wall motion may be an important feature in some of these systems. For instance, chondrocytes express K_ir_ channels^36^, gap junctions^37^, and the ionotropic serotonin receptor 5-ht3a^38^, suggesting that the ingredients are present for regulation of membrane potential via domain wall migration.

The shape of the *I*-*V* curve in our experiments is qualitatively captured by a cubic (Fitzhugh Nagumo-type) nonlinearity.^39, 40^ Models of this sort have been applied to describe similar dynamics (zero-dimensional hysteresis, domain nucleation, growth and disorder-driven breakup) in magnetic domain reversals in ferromagnets,^41^ in the spread of forest fires,^42^ in phase transitions,^26^ and in expanding species ranges with a strong Allee effect.^43^ Our work shows that the reaction-diffusion formalism can also be applied to spatial symmetry breaking in electrophysiology.

In the Turing model, formation of quasi-periodic patterns requires interaction of two or more morphogens, often described as an activator and an inhibitor. The Hodgkin-Huxley equations describe the dynamics of voltage, a scalar quantity. The spatial symmetry breaking studied in this report does not constitute a classical Turing pattern, in that voltage is only a one-dimensional state variable. As a result, the patterns of membrane voltage did not have a characteristic finite spatial frequency. To achieve a classical Turing-like pattern would require coupling of voltage to another diffusible species, e.g. Ca^2+^. It is not known whether classical Turing-like patterns of membrane voltage can be created.

## Acknowledgments

We thank Urs Boehm, Amanda Klaeger, Juanita Mathews, and Michael Levin for helpful discussions. We thank Evan Miller for help providing the BeRST1 dye.

## Funding

This work was supported by the Allen Discovery Center at Tufts University, the Vannevar Bush Fellowship Foundation, and the Howard Hughes Medical Institute. HMM was supported by the Department of Defense (DoD) through the National Defense Science & Engineering Graduate Fellowship (NDSEG) Program. GO was supported by the Howard Hughes Medical Institute Gilliam Fellowship.

## Author Contributions

HMM and AEC conceived and designed the study. HMM conducted and analyzed experimental results, with assistance from RJS, HX, and SB. HMM designed and simulated numerical models of electrically bistable cells. ZAT and OP provided hiPSC-derived myoblasts for myocyte differentiation and characterized these cells via immunocytochemistry and RNAseq. GO provided BeRST1 dye reagent. HMM and AEC wrote the manuscript, with input from ZAT and OP. AEC and OP oversaw the research.

## Competing Interests

The authors declare no competing financial interests.

## Data and Materials Availability

All data are available in the main text or the supplementary materials. Raw data are available from the corresponding author upon reasonable request. The materials that support the findings of this study are available from the corresponding author upon reasonable request.

## Methods

### Experimental design

The goal of this study was to discover and characterize bioelectrical domain walls: electrophysiological entities through which tissues can mediate long-range electrical signaling without using action potentials. We modeled electrically bistable cells as a one-component reaction-diffusion system with a bistable electrical nonlinearity. The nonlinear *I-V* curve was composed of an inward-rectifying potassium conductance (K_ir_2.1) and an Ohmic leak conductance with reversal potential near 0 mV.

To explore bioelectrical domain walls experimentally, we generated an engineered electrically bistable HEK cell line (bi-HEKs) which expressed K_ir_2.1 and light-gated channelrhodopsin leak conductance. To probe for domain wall formation in a physiological system we studied the electrophysiology of myocytes during differentiation. We chose this system because myoblast precursors gradually express an inward rectifying potassium conductance during differentiation, and myocyte differentiation requires spatial coupling between myoblast precursors. Myogenesis therefore offered a plausible physiological system in which to observe bioelectrical domain wall formation.

### Numerical modeling of electrically bistable cells and tissues

In the conductance-based model, the voltage dynamics are governed by the equation:

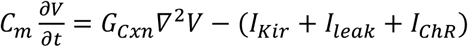

The gap junction sheet conductance is defined as *G_Cxn_* = *g_Cxn_* × *l*^2^, where *g_Cxn_* is the gap junction conductance between adjacent cells, and *l* is the linear dimension of a cell. If *g_Cxn_* is measured as an areal conductance (S/m^2^), then the units of *G_Cxn_* are *S* (i.e. sheet conductance, which has no spatial dimension). If *C*_m_ is measured as an areal capacitance (F/m^2^), then the ratio *G_Cxn_/C_m_* has units of a diffusion coefficient (m^2^/s), making explicit the connection between the Hodgkin-Huxley equations and the reaction-diffusion equation.

For convenience in simulations, units of space were 10 μm (corresponding to linear size of one cell), units of time were ms, and units of voltage were mV. We assumed the capacitance of a cell was 10 pF. Conductances were measured in nS/pF, ionic currents in pA/pF, and voltage in mV. Parameters used in all simulations are given in the Tables in the *Model parameters* section of the *Methods*.

Simulations were run in Matlab using custom software. Single-cell voltages in 0D were determined by finding fixed points of cell-autonomous current-voltage curves. Extended tissues were numerically simulated as two-dimensional grids of 100 x 100 cells with periodic boundary conditions and one grid-point per cell. Simulations and experiments thus had similar spatial scales. For simulations of nucleation events in homogenous tissues (Fig 3B), 300 x 300 cell grids with no-flux boundary conditions were implemented to match experimental conditions. The discrete Laplacian was implemented using the matlab `del2` function (with default spacing) and solutions were time-integrated using Euler’s method with 10 kHz sampling.

The inward rectifying potassium current from K_ir_2.1 was based on a model from ten Tusscher *et al.* ^21^ as: *I*_*K*_ = *g*_*K*_*x*_*K*∞_(*V*)(*V* − *E*_*K*_) with reversal potential at EK = −90 mV. The parameter *x*_*K*∞_(*V*) is a time-independent rectification factor that depends on voltage, with the following form:

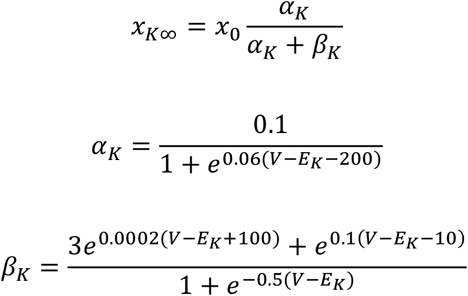

The scaling factor *x*_0_ = 100 was introduced to make *x*_*K*∞_ of order unity between −90 and −60 mV. In our simulations the conductance magnitude *g*_*K*_ was the only parameter varied to mimic changes in expression of K_ir_2.1. The variable leak was modeled as an Ohmic conductance with reversal potential 0 mV. For homogeneous tissues all cells had identical K_ir_2.1 and leak conductances.

To introduce disorder into the tissue, a fraction *n*_K_ of the cells were randomly assigned to express K_ir_2.1 (all with conductance *g_K_*) while the remaining cells had no K_ir_2.1 expression (Figure S2, S3). Spatial correlations in K_ir_2.1 expression were introduced by assigning a random number to each cell, independently sampled from a uniform distribution on [0, 1]. The values where then smoothed with a two-dimensional Gaussian kernel of width *d*. A threshold was selected so that a fraction *n*_K_ of the cells were above threshold. These cells were assigned to express K_ir_2.1 with conductance *g_K_* and cells below threshold did not express. The extent of the spatial correlations was tuned by varying *d*. In the simulation for Fig. 3h, K_ir_2.1 and CheRiff expression were heterogeneous, and independent of each other. The distribution of CheRiff expression was calculated following the same procedure as for K_ir_2.1, using the same smoothing parameter, *d*, but different thresholds *n*_K_, and *n_ChR_*.

Cells in simulated tissues were first initialized to their cell-autonomous resting potential, i.e. the resting potential in the absence of influences from the neighbors. Bistable cells were initialized to the fixed point which had the greater area under the curve between it and the unstable fixed point. Tissues were then time-evolved to generate an initial steady state voltage profile. To simulate bioelectrical dynamics under changes in parameters, conductances were gradually changed over timescales slower than any of the internal relaxation dynamics. For simulations of domain boundary velocities in homogeneous tissues with bistable *I-V* curves (Fig. S1), the left half of the tissue was initialized in a depolarized state and the right half in a hyperpolarized state.

### Bi-HEK cell generation and culture

Genetic constructs encoding the inward rectifying potassium channel K_ir_2.1 and the blue-shifted channelrhodopsin CheRiff were separately cloned into lentiviral expression backbones (FCK-CMV) and then co-expressed in HEK 293T cells along with the lentiviral packaging plasmid PsPAX2 (Addgene) and the envelope plasmid VsVg (Addgene) via polyethylenimine transfection (Sigma). Lentiviral particles were harvest at 36 and 72 hours post-transfection, and then concentrated 20-fold using the Lenti-X concentrator system (Takara).

For experiments where nominally homogeneous expression was the goal (Figs. 2, 3), HEK cells were incubated with both K_ir_2.1 and CheRiff lentiviral vectors for 48 hours prior to measurement, and then passaged and replated onto poly-D-lysine coated glass-bottom tissue culture dishes (MatTek). For patch clamp measurements (Fig. 2), bi-HEKs were plated sparsely onto Matrigel coated dishes. For wide-field measurements (Fig. 3), adhesive islands were prepared by manually spotting poly-d-lysine onto MatTek plates. Bi-HEKs were plated onto these plates to create confluent patches of cells approximately 2 mm in diameter.

For experiments where disordered expression was the goal (Supplementary Fig. 3), K_ir_2.1 and CheRiff constructs were transiently expressed (using Mirus 293T) in HEK cells which were then grown to confluence for an additional 72 hours prior to measurement. High-disorder samples were not replated prior to measurement.

### hiPSC myoblast and myocyte differentiation

HiPSC-derived myoblasts were differentiated into myocyte fibers according to an established serum-free differentiation protocol.^44^ Briefly, 3-4 weeks old primary myogenic cultures generated from wild-type hiPSCs were dissociated as described and myogenic progenitors (myoblasts) were replated at low density (35-40k/cm^2^) onto Matrigel (Corning, Cat#354277)-coated dishes in skeletal muscle growth media (SKGM-2, Lonza CC-3245) with 10 μM ROCK inhibitor.^44^ After 24 hours, medium was changed to SKGM-2 media without ROCK inhibitor and incubated with low-titer lentivirus encoding CheRiff-CFP. Myoblast cultures were proliferated for up to 72 hours, at which point cultures reached ∼90% confluence. Cultures were then induced for myogenic differentiation with DMEM/F12 supplemented with 2% knock-out serum replacement (Invitrogen, Cat. # 10828028), 10 μM of the TGFβ inhibitor SB431542 (Tocris, Cat. # 1614), 1 μM Chiron (Tocris, Cat. # 4423), 0.2% Pen/Strep (Life Technologies, Cat. # 15140122) and 1x ITS (Life Technologies, Cat. # 41400045). ^44^ Following induction, medium was changed on days 1 and 2 and then was refreshed every other day for up to 10 days post-differentiation to generate mature and fused myocyte fibers. Samples were measured between 0 and 3 days after start of differentiation (i.e., after 3 and 6 days in culture).

### Immunostaining and imaging

Human iPSC-derived myocyte cultures were fixed for 20 minutes in 4% formaldehyde. Cultures were rinsed three times in phosphate-buffered saline (PBS), followed by blocking buffer composed of PBS supplemented with 10% fetal bovine serum (FBS) and 0.1% Triton X-100. Primary antibodies were then diluted in blocking buffer and incubated overnight at 4 °C. Cultures were then washed three times with PBST (PBS supplemented with 0.5% Tween-20) and incubated with secondary antibodies conjugated with an AlexaFluor dye (Molecular probes) and DAPI (5 μg/mL) in blocking buffer for 2 h at room temperature. Cultures were washed with PBST followed by PBS, followed by imaging. Antibodies were: anti-PAX7 (Developmental Studies Hybridoma Bank, DSHB), anti-Myogenin (Santa Cruz, SC-576X) and embryonic anti-MyHC (DSHB, F1.652).

### Transcriptomic profiling: Library preparation and sequencing

RNA was extracted from cells using Trizol (Invitrogen) or with the RNeasy Mini Kit (Qiagen). Libraries were prepared using Roche Kapa mRNA HyperPrep sample preparation kits from 100 ng of purified total RNA according to the manufacturer’s protocol. The finished dsDNA libraries were quantified by Qubit fluorometer, Agilent TapeStation 2200, and RT-qPCR using the Kapa Biosystems library quantification kit according to manufacturer’s protocols. Uniquely indexed libraries were pooled in equimolar ratios and sequenced on two Illumina NextSeq500 runs with single-end 75bp reads by the Dana-Farber Cancer Institute Molecular Biology Core Facilities.

### Transcriptomic profiling: RNAseq Analysis

Sequenced reads were aligned to the UCSC hg19 reference genome assembly and gene counts were quantified using STAR (v2.5.1b)^45^. Differential gene expression testing was performed by DESeq2 (v1.10.1)^46^ and normalized read counts (FPKM) were calculated using cufflinks (v2.2.1)^47^. RNAseq analysis was performed using the VIPER snakemake pipeline.^48^

### Patch clamp and all-optical electrophysiology

All electrophysiological measurements were performed in Tyrode’s solution, containing (in mM) 125 NaCl, 2 KCl, 2 CaCl_2_, 1 MgCl_2_, 10 HEPES, 30 glucose. The pH was adjusted to 7.3 with NaOH and the osmolality was adjusted to 305–310 mOsm with sucrose. Prior to measurements, 35-mm dishes were washed twice with 1 mL phosphate-buffered saline (PBS) to remove residual culture media, then filled with 2 mL Tyrode’s solution.

For patch clamp measurements, filamented glass micropipettes (WPI) were pulled to a resistance of 5– 10 MΩ and filled with internal solution containing (in mM) 140 KCl, 1 MgCl_2_, 10 EGTA, 10 HEPES, 3 Mg-ATP, pH adjusted to 7.3 with KOH. The patch electrode was controlled via a low-noise patch clamp amplifier (A-M Systems, model 2400). Voltage traces were collected under *I* = 0 current clamp mode, and current traces were collected in voltage clamp mode. Blue light for optical stimulation (Coherent Obis 488 nm) was modulated using an acousto-optic tunable filter (Gooch and Housego GH18A series). Patch clamp measurements were performed on small clusters of cells (approximately 6 cells).

Spatially resolved optical electrophysiology measurements were performed using a home-built upright ultra-widefield microscope^49^ with a large field of view (4.6×4.6 mm^2^, with 2.25×2.25 μm^2^ pixel size) and high numerical aperture objective lens (Olympus MVPLAPO 2XC, NA 0.5). Fluorescence of BeRST1 was excited with a 639 nm laser (OptoEngine MLL-FN-639) at 100 mW/cm^2^, illuminating the sample from below at an oblique angle to minimize background autofluorescence. BeRST1 fluorescence was separated from scattered laser excitation via a dichroic beam splitter (Semrock Di01-R405/488/561/635-t3-60×85) and an emission filter (Semrock FF01-708/75-60-D). Images were collected at 100 Hz frame rate on a Hamamatsu Orca Flash 4.2 scientific CMOS camera. Optogenetic stimulation was performed by exciting CheRiff with a blue LED (Thorlabs M470L3) with a maximum intensity of at 10 mW/cm^2^.

Prior to measurement, cells were incubated with 1 μM BERST1 dye in phosphate buffered saline for 30 minutes in a tissue culture incubator. Samples were then washed and prepared in Tyrode’s solution immediately before imaging.

### Data Analysis and Image Processing

Optical recordings of voltage-sensitive BeRST1 fluorescence were acquired for isolated cells, small clusters of cells (4-12 cells), and extended tissues (>2 mm linear size). Recordings were processed using custom MATLAB software. Briefly, to minimize uncorrelated shot-noise, movies were subjected to 4×4 binning, followed by pixel-by-pixel median filtering in the time domain (9 frame kernel). A background signal was calculated from a cell-free region of the field of view and subtracted from the region containing the cells. Mean sample images were generated by measuring the average fluorescence of the tissue prior to optogenetic stimulation. Functional recordings were divided pixel-wise by this baseline to generate movies of ΔF/F. Plots of voltage-dependent fluorescence were generated by averaging the time-lapse movies over the relevant region of interest (e.g. small clusters for 0D data; localized spots within extended cell culture islands for 2D local measurements; and over entire cell culture islands for 2D mean measurements).

### Statistical Analysis

Statistical analysis was performed on optical electrophysiology recordings from immature myocyte precursors to assess the significance of population-level differences in electrophysiological phenotypes at different time points. Measurements were performed on populations of isolated cells taken at 3 and 6 days in culture (see hiPSC myoblast and myocyte differentiation methods above). Fluorescent recordings of voltage were acquired for 100 ms prior to illumination, during a 40 s ramp of blue light (increasing from 0 to 10 mW/cm^2^ for 20 s, then decreasing for 20 s), and then for 5 s after the blue light was turned off. Raw acquisitions were converted to ΔF/F for further analysis.

Electrophysiological phenotypes were parameterized by calculating the total endpoint hysteresis and the mean integral hysteresis for each identified cell (Supplementary Fig. 5). Endpoint hysteresis was defined as the difference in ΔF/F between the 5 s at the end of the acquisition and the 100 ms at the start of the acquisition. Integral hysteresis was defined as the difference between the mean ΔF/F over the decreasing phase of the blue light ramp and the mean ΔF/F during the increasing phase of the blue light ramp (each averaged over the corresponding full 20 s ramp).

In each sample, individual cells which responded to CheRiff stimulation were first manually identified via an overall mean ΔF/F image. 51 and 89 cells were identified in the day 3 and day 6 measurements, respectively. Endpoint hysteresis and mean hysteresis were then calculated for each identified cell, and a cell was sorted as hysteresis-positive if it demonstrated a value greater than 0.3 for either measure (Figure S4). Cells which showed endpoint hysteresis > 0.3 were sorted into the endpoint hysteresis cluster, regardless of their integral hysteresis value. Cells with integral hysteresis > 0.3 and endpoint hysteresis < 0.3 were sorted into the integral hysteresis cluster. Statistical errors were estimated as the standard deviation from a binomial distribution; i.e.,

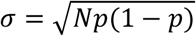

The significance of population level phenotypes was assessed via paired two-sample t-tests using standard MATLAB functions (ttest2). Smooth depolarizations without hysteresis were observed on day 3 in 34 of 51 cells and on day 6 in 42 of 89 cells, a significant decrease (p = 0.026). Endpoint hysteresis was observed on day 3 in 17 of 51 cells and on day 6 in 26 of 89, not a significant difference (p = 0.61). Integral hysteresis was observed on day 3 in 0 of 51 cells and on day 6 in 21 of 89 cells (p <0.001).

## Supplementary Information

**Tables.** Model parameters

| Parameter | Explanation |
| --- | --- |
| $C_m = 10 \text{ pF}$ | Single cell capacitance |
| $l = 10 \text{ }\mu\text{m}$ | Size of an individual cell; discretization size of simulated tissues |
| $E_K = -90 \text{ mV}$ | Potassium reversal potential |
| $E_{ChR} = 0 \text{ mV}$ | Channelrhodopsin reversal potential |
| $E_{leak} = -20 \text{ mV}$ | Additional leakage reversal potential. Used for myocyte models (Fig. 4) and for domain model (Fig. S3). |
| $G_{Cxn} = 2 \text{ nS/pF} \times l^2$<br>$= 200 \text{ (nS/pF)} \times \mu\text{m}^2$ | Strength of gap junction coupling between bi-HEK cells |
| $n_{Kir} = 0 - 1$ | Proportion of cells initialized to express Kir2.1 |
| $n_{ChR} = 0 - 1$ | Proportion of cells initialized to express CheRiff. Expression is uniform ($n_{ChR} = 1$) in all panels except for Fig. 3h. |
| $d = 0.001 \text{ to } 10 \text{ cells}$ | Width of gaussian smoothing kernel used to introduce spatial correlations in Kir2.1 expression |
| $g_K = 0 - 500$ pS/pF | Kir2.1 conductance. This conductance is multiplied by the inward rectifying function $x_{K\infty}$. |
| $x_0 = 100$ | Scaling factor to set $x_{K\infty}$ to order unity between $E_K$ and the point of maximum outward current ($V_{max} \sim -69$ mV) |
| $g_{ChR} = 0$ to 100 pS/pF | Conductance of the channelrhodopsin CheRiff. Varied continuously in simulations of blue light intensity ramps. |
| $g_{leak} = 5$ pS/pF | Additional leak conductance added to myocyte model |

Model parameters for figure panels:
| Figure panel | Parameters |
| --- | --- |
| 2B | $g_{ChR} = 50$ pS/pF, $g_K = 200$ pS/pF |
| 3A | $g_{ChR} = 50$ pS/pF, $g_K = 200$ pS/pF, $n_{Kir} = 1$ |
| 3B, top | $g_{ChR} = 70$ pS/pF, $g_K = 200$ pS/pF, $n_{Kir} = 1$ |
| 3B, bottom | $g_{ChR} = 60$ pS/pF, $g_K = 200$ pS/pF, $n_{Kir} = 1$ |
| 3C | $g_{ChR} = 1$ to 200 pS/pF, $g_K = 200$ pS/pF, $n_{Kir} = 1$ |
| 4E | $g_{leak} = 100$ pS/pF, $g_{ChR} = 0.1$ to 300 pS/pF (ramped),<br>$g_K = 120, 240, 480$ pS/pF (left, middle, and right panels) |
| S2B | $g_{ChR} = 24.5$ pS/pF, $g_K = 60, 72.5, 85, 97.5$ pS/pF (top to bottom) |
| S2C | $g_{ChR} = 22$ pS/pF, $n_{Kir} = 1$ |
| S3B | $g_K = 1 \text{ to } 500 \text{ pS/pF}$, $g_{ChR} = 20 \text{ pS/pF}$, $n_{Kir} = 1$ |
| S3D,E | $g_K = 500 \text{ pS/pF}$, $g_{ChR} = 20 \text{ pS/pF}$, $d = 0.001$, $n_{Kir} = 0 \text{ to } 1$ |
| S3G,H | $g_K = 500 \text{ pS/pF}$, $g_{ChR} = 20 \text{ pS/pF}$, $d = 10$, $n_{Kir} = 0 \text{ to } 1$ |
| S4C,D | $g_K = 50 \text{ pS/pF}$, $g_{leak} = 20 \text{ pS/pF}$, $E_{leak} = -20 \text{ mV}$, $d = 3$, $n_{Kir} = 0.25$, $g_{ChR} = 0 \text{ to } 250 \text{ pS/pF}$ |
| S4G,H | $g_{ChR} = 1 \text{ to } 300 \text{ pS/pF}$, $g_K = 600 \text{ pS/pF}$, $n_{Kir} = 0.5$, $n_{ChR} = 0.75$, $d = 1.5$ |
| S5A | $g_{leak} = 100 \text{ pS/pF}$, $g_{ChR} = 0.1 \text{ to } 300 \text{ pS/pF}$ (ramped),<br>$g_K = 120, 240, 480 \text{ pS/pF}$ (left, middle, and right panels) |

## Supplementary Figures

**Supplementary Figure 1.**
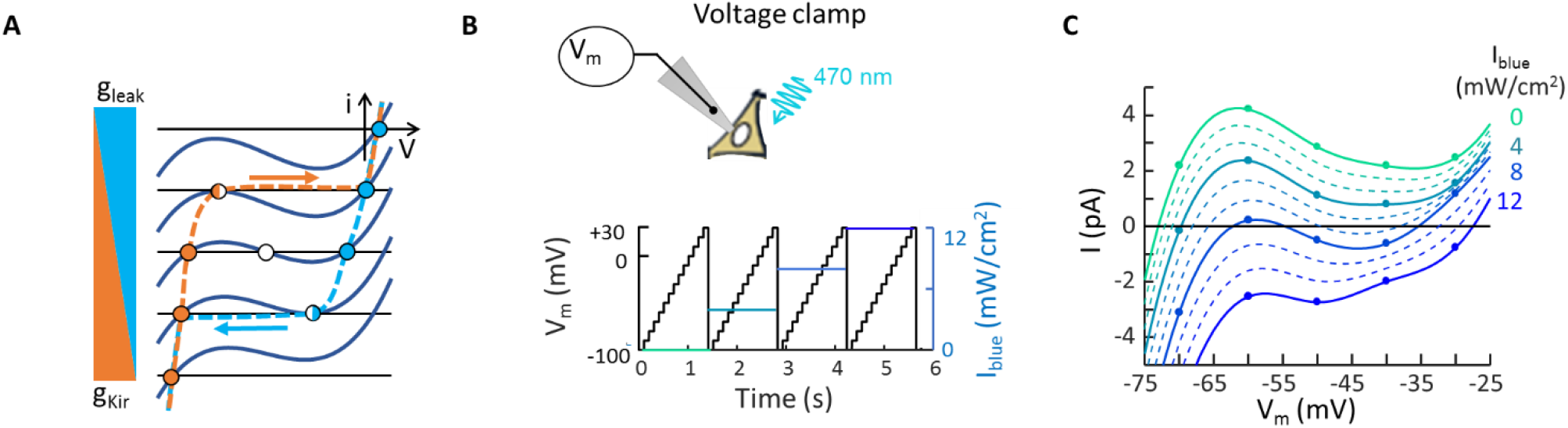
Electrophysiological characterization of bi-HEK cells. A) Cartoon showing origin of bistability in cells expressing a K_ir_ channel and a leak conductance. Changes in the ratio of leak to K_ir_ conductance drive the *I*-*V* curve through two saddle-node bifurcations. B) Protocol for measuring the *I*-*V* curve under different levels of optogenetic drive. Measurements were performed in voltage clamp mode. C) Patch clamp measurements of the *I*-*V* curve of a small island of bi-HEKs under varying blue light illumination. Points represent measurements. Dotted lines are interpolations.

**Supplementary Figure 2.**
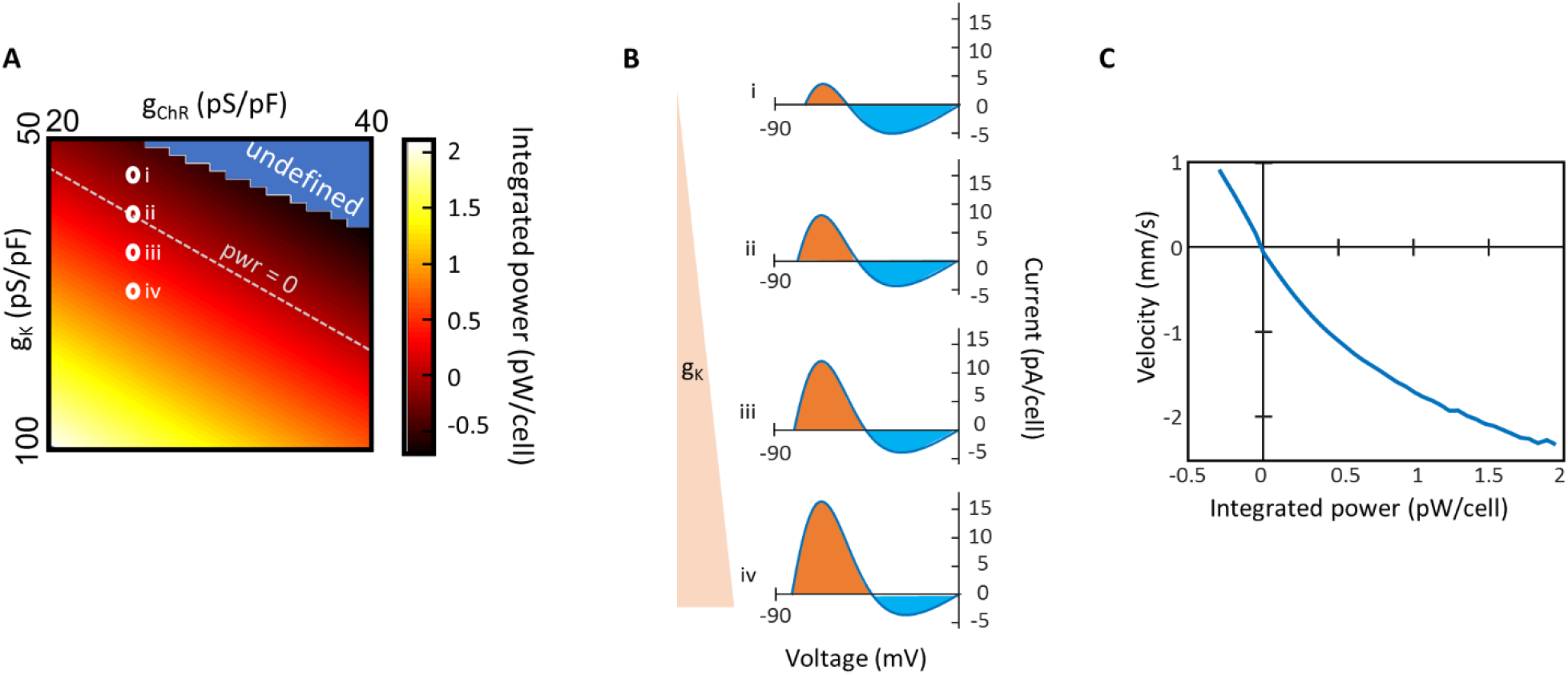
Balance of ionic currents determines velocity of domain wall motion in homogeneous tissues. The balance of currents is set by the integral of the *I-V* curve between the two stable fixed points, or equivalently the relative areas of the orange and blue shaded regions in panel (B). The area under the curve (∫ *IdV*) has units of power. A) Area under the *I-V* curve as a function of the K_ir_2.1 and channelrhodopsin conductance levels. The white dotted line corresponds to a stationary domain boundary. In the blue region there is only one stable fixed point (near V = 0), so the area under the curve is undefined. B) Example *I-V* curves from the corresponding circled regions in (A). C) Domain wall velocity as a function of the area under the *I*-*V* curve. Positive velocities indicate growth of the depolarized domain.

**Supplementary Figure 3.**
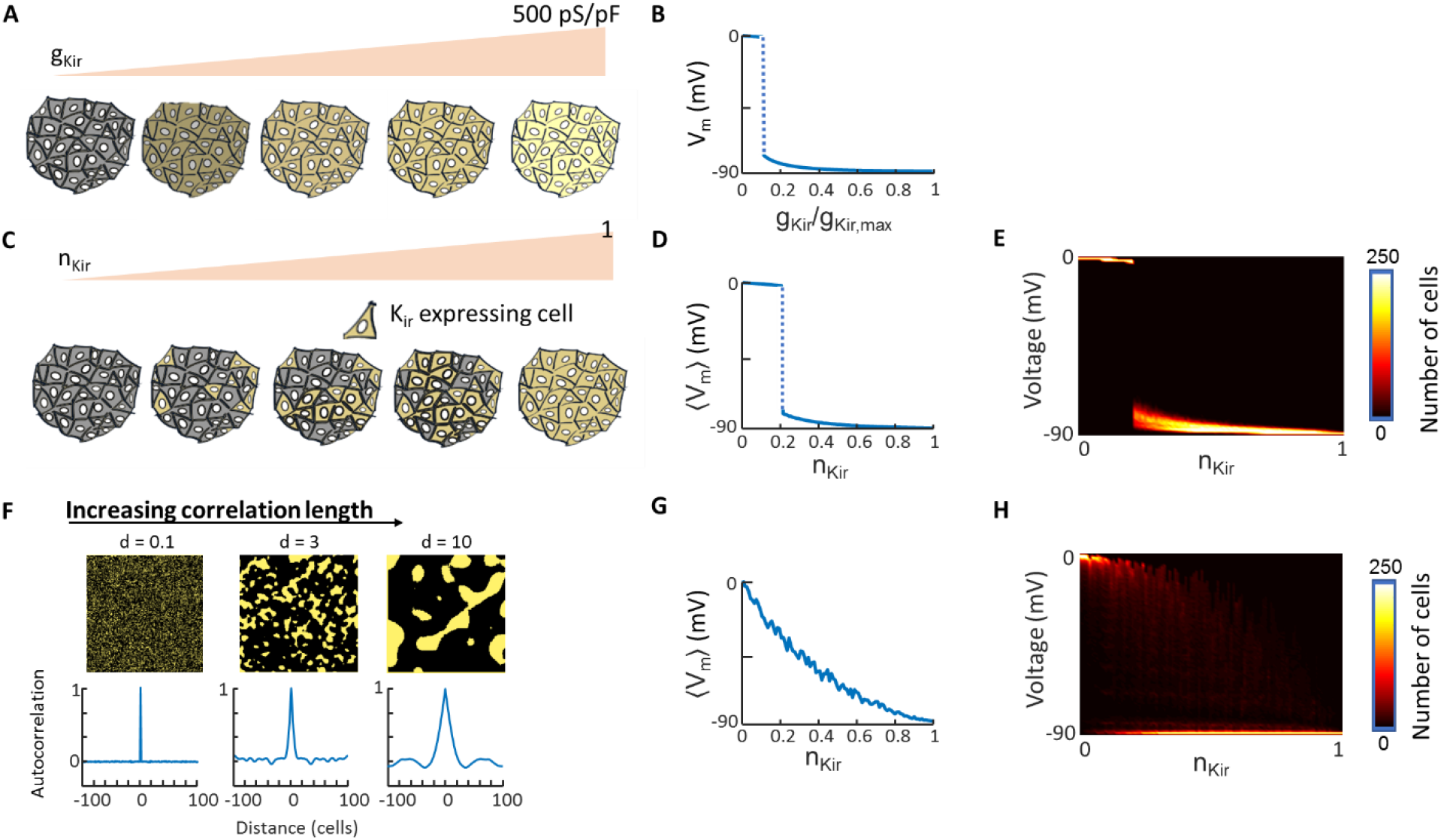
Effect of noisy gene expression on domain wall properties. A) Cartoon showing a homogeneous tissue with constant leak conductance and gradually increasing K_ir_2.1 conductance. B) Membrane voltage (Vm) as a function of *g*_Kir_. Abrupt polarizing transition arises when *g*_Kir_ is sufficient to drive polarized domain wall growth. C) Cartoon showing a heterogeneous tissue where the fraction of cells that express K_ir_2.1 gradually increases. In this cartoon the probability of expression is independent in each cell. D) As in the homogeneous tissue, at a critical expression density the tissue-average voltage polarizes in a step-wise manner due to domain wall migration. E) Probability distribution of single-cell voltage values as a function of *n*_Kir_. In the heterogeneous tissue, the voltage can vary between cells, though when the size of each cell is much smaller than the domain wall width, 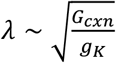, then the *g*_*K*_ distribution of single-cell voltages is narrowly centered around the mean. F) Introduction of spatially correlated gene expression in models of bi-HEK cells. Top: example images of K_ir_2.1 expression with different degrees of spatial correlation. Bottom: radially averaged autocorrelation functions of the simulated tissues shown above. G) Tissue-average membrane potential as a function of *n*_Kir_ in the case of correlated disorder (*d* = 10 cells). The step-wise transition to polarization is replaced by a patchwork of polarized and depolarized domains. Changes in *n*_Kir_ change the relative populations of these two domains. H) Probability distribution of single-cell voltage values as a function of *n*_Kir_ in the tissue simulated in (G).

**Supplementary Figure 4.**
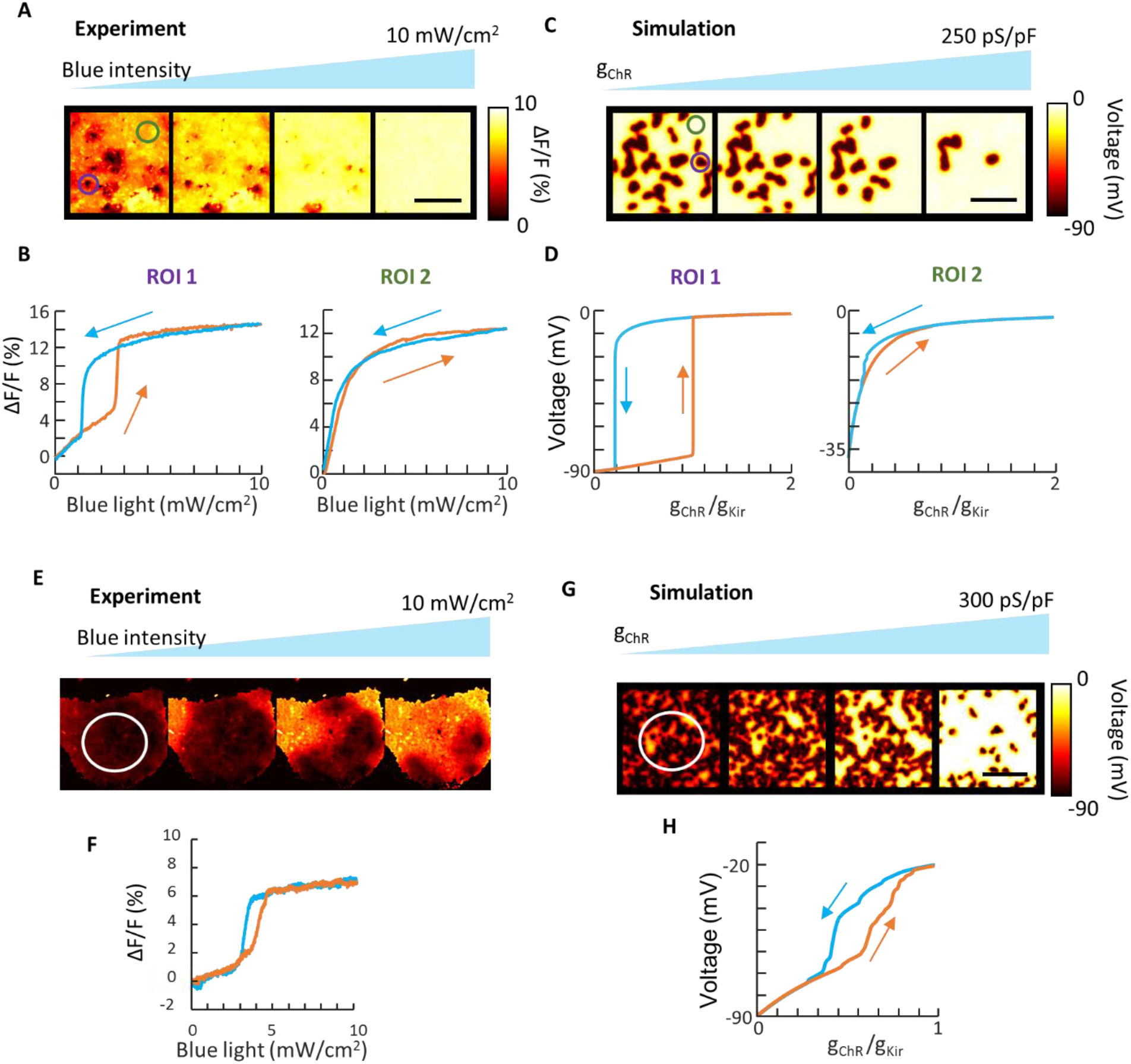
Spatially correlated variability in K_ir_2.1 expression drives breakup of tissues into discrete domains. A monoclonal cell line expressing CheRiff only was transfected with plasmid for K_ir_2.1 and then allowed to grow for 3 days to reach confluence, leading to clusters of correlated expression ∼3 cells wide. Calibration transfections with a fluorescent marker showed that only ∼30% of the cells expressed. A) Images of fluorescence increase (ΔF/F) in a heterogeneous culture under ramped optogenetic stimulation. Scale bar 1 mm. B) Plots of fluorescence from the two indicated regions of interest (ROIs) in (A). Some ROIs showed hysteresis while others showed smooth and reversible depolarization. C) Frames from a simulation with noisy K_ir_2.1 expression and ramped optogenetic drive. Scale bar 0.5 mm. D) Regions of the simulation showed ROIs with hysteresis and others without. E) Island of bi-HEK cells driven with a triangle wave of blue light showing depolarization via domain wall migration. F) Fluorescence averaged over the circled region in (E), showing partial hysteresis and Barkhausen-like noise in a disordered sample. G) Simulation of depolarization of a noisy tissue under ramped optogenetic stimulation. H) Mean voltage in the circled region in (G) showing partial hysteresis and Barkhausen-like noise in a disordered sample.

**Supplementary Figure 5.**
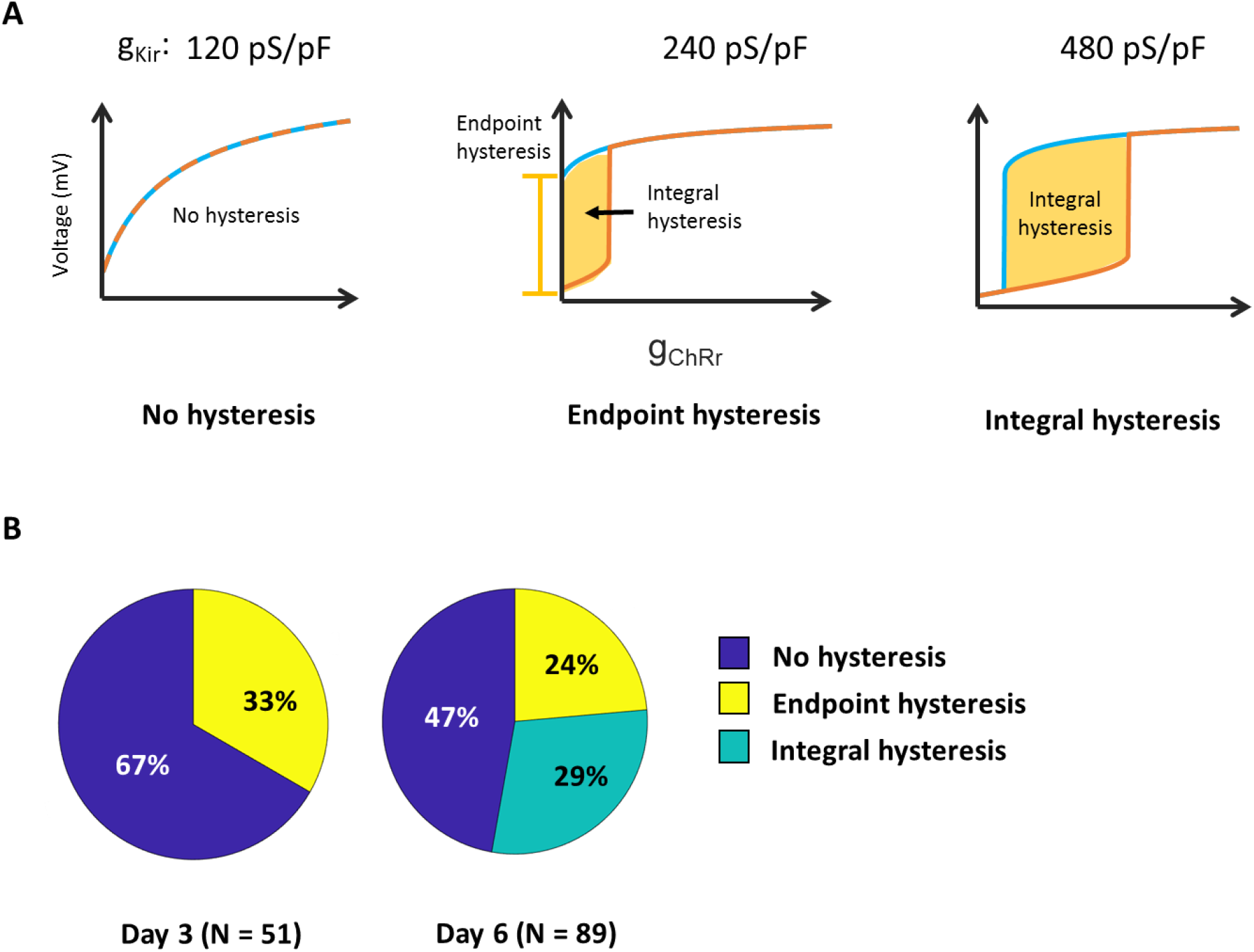
Electrophysiological phenotypes in immature isolated myocytes. A) Definition of electrophysiological classes showing “No hysteresis”, “Endpoint hysteresis” and “Integral hysteresis”. The plots show numerical simulations of the K_ir_ + leak + ChR model with increasing levels of K_ir_ and all other parameters held constant. B) Distribution of phenotypes by day of measurement.

**Supplementary Figure 6.**
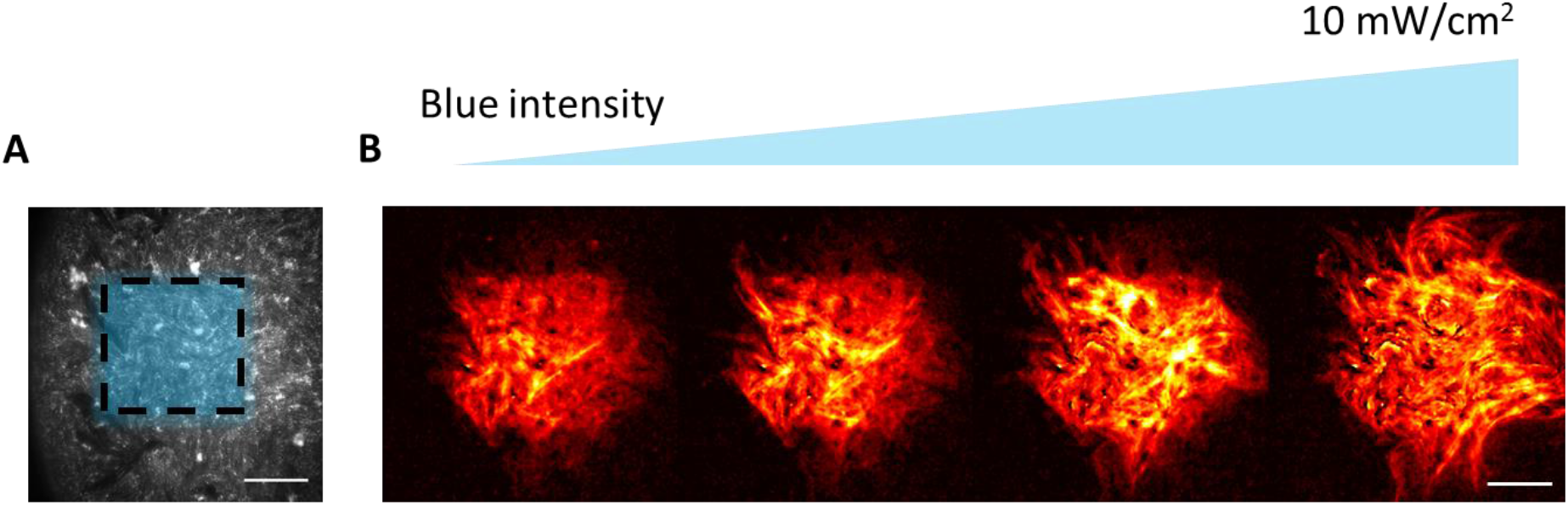
Gap junction coupling in confluent cultures of immature myocytes. A) image of the myocyte culture showing region of cells stimulated with blue light. B) Fluorescence of BeRST1 indicating electrical depolarization as a function of optogenetic stimulus strength. Propagation of the depolarization to cells outside the stimulated region establishes the presence of gap junction-mediated electrical coupling in the culture. Scale bars 1 mm.

## Supplementary Movie Captions

**Supplementary Movie S1: Simulation of nucleation and growth of bioelectrical domains in homogeneous tissue**

**Supplementary Movie S2: Nucleation and growth of bioelectrical domains in a confluent culture of bi-HEK cells, example 1**

**Supplementary Movie S3: Nucleation and growth of bioelectrical domains in a confluent culture of bi-HEK cells, example 2**

**Supplementary Movie S4: Switching of discrete bistable bioelectrical domains in disordered culture of bi-HEK cells**

**Supplementary Movie S5: Simulation of switching of discrete bistable bioelectrical domains in disordered tissue**

**Supplementary Movie S6: Depolarization via domain wall propagation in a confluent culture of human iPSC-derived myocytes**

## Data Availability

All data supporting the conclusions are available from the authors on reasonable request.

## Code Availability

Custom-written code used for data analysis is available from the authors on request.

## Material Availability

Lentiviral expression plasmids for CheRiff and K_ir_2.1 are available from the authors upon reasonable request.

## Notes

#### Summary of Updates

Revisions to Figs.

